# Single-Cell Antigen Receptor Sequencing in Pigs with Influenza

**DOI:** 10.1101/2024.10.13.617920

**Authors:** Weihong Gu, Darling Melany de Carvahlo Madrid, Sadie Clements, Laurie Touchard, Nathan Bivins, Grant Zane, Mingyi Zhou, Kiho Lee, John P. Driver

## Abstract

Understanding the pulmonary adaptive immune system of pigs is important as respiratory pathogens present a major challenge for swine producers and pigs are increasingly used to model human pulmonary diseases. Single-cell RNA sequencing (scRNAseq) has accelerated the characterization of cellular phenotypes in the pig respiratory tract under both healthy and diseased conditions. However, combining scRNAseq with recovery of paired T cell receptor (TCR) α and β chains as well as B cell receptor (BCR) heavy and light chains to interrogate their repertoires has not to our knowledge been demonstrated for pigs. Here, we developed primers to enrich porcine TCR α and β chains along with BCR κ and λ light chains and IgM, IgA, and IgG heavy chains that are compatible with the 10x Genomics VDJ sequencing protocol. Using these pig-specific assays, we sequenced the T and B cell receptors of cryopreserved lung cells from *CD1D*-expressing and -deficient pigs after one or two infections with influenza A virus (IAV) to examine whether natural killer T (NKT) cells alter pulmonary TCR and BCR repertoire selection. We also performed paired single-cell RNA and receptor sequencing of FACS-sorted T cells longitudinally sampled from the lungs of IAV-vaccinated and -infected pigs to track clonal expansion in response to IAV exposure. All pigs presented highly diverse repertoires. Pigs re-exposed to influenza antigens from either vaccination or infection exhibited higher numbers of expanded CD4 and CD8 T cell clonotypes with activated phenotypes, suggesting potential IAV reactive T cell populations. Our results demonstrate the utility of high throughput single-cell TCR and BCR sequencing in pigs.

## Introduction

High throughput single-cell RNA sequencing (scRNAseq) technology has greatly increased our understanding of the phenotypic diversity and plasticity of immune cell types as well as their cellular interactions in complex tissues in both healthy and disease states (1, 2). In addition to revealing cellular heterogeneity through RNA expression, scRNAseq can be coupled with additional assays to enhance cellular phenotyping, such as enrichment of T cell receptor (TCR) and B cell receptor (BCR) repertoires using primers that target the C regions in mRNA transcripts of TCR and BCR chains and isotypes. This allows construction of expressed TCRs and BCRs of individual cells including genomic rearrangements between the variable (V), diversity (D), and joining (J) regions of the TCR and BCR intervals responsible for creating diversity in receptor binding surfaces. Pairing TCR/BCR sequencing with scRNAseq provides a powerful approach for studying the relationship between immune repertoire and many types of immune responses. Additionally, V(D)J recombination at the TCR and BCR loci can be used as endogenous barcodes to trace T and B cell clonotypes as they expand or transition though different states, including within the same individual over time (3, 4).

Domestic pigs (*Sus scrofa*) are an important agricultural species that are intensively farmed making them vulnerable to many infectious pathogens. Therefore, a thorough understanding of the porcine immune system is needed to optimize vaccine and drug design and to identify immune targets for increased disease resistance through selective breeding and genetic engineering. Because of their many similarities to humans, swine are increasingly used in place of non-human primate models (5, 6). However, a limitation that prevents fully exploiting these pig models is our incomplete understanding of the porcine immune system which is due in part to a scarcity of immune profiling reagents for pigs. scRNAseq which does not require marker-based sorting of cell subsets is helping to address this gap.

In the current work, we developed porcine-specific TCR and BCR primers compatible with the droplet-based protocols of the 10x Genomics Next GEM Single Cell 5’ sequencing protocol. These assays were used to compare cryopreserved lung cells from pigs genetically engineered to lack *CD1D*, which encodes an antigen presenting molecule required for the development of natural killer T (NKT) cells, a subset of innate-like T cell that accumulates in barrier organs such as the lungs (7–12). Pigs were analyzed after one or two infections with influenza A virus (IAV) to interrogate the evolution of the TCR and BCR repertoires after primary or secondary infection and to determine if NKT cells exert T helper cell functions that influence receptor diversity. Additionally, we applied our protocol to a longitudinal assessment of T cells recovered from the lung lavage fluid of infant pigs exposed to IAV vaccination and infection. Here we were able to track clonal expansion and monitor changes in the frequency of clonotypes within the same pig. Together, these results demonstrate the potential of TCR/BCR profiling to better understand a wide variety T and B cell-related immune responses in pigs.

## Results

### Single-cell RNA sequencing analysis of influenza virus-infected genetically edited pigs

Cryopreserved cells liberated from the enzyme digested lungs of mixed-breed pigs carrying an inactive form of the *CD1D* (*CD1D−/−*) gene compared with littermates carrying one copy (*CD1D−/+*) were subjected to paired scRNAseq and scTCR/BCRseq using our custom primer sets. One cohort of pigs was necropsied five days after a single infection (1X) with H1N1 A/Missouri/CS20N08/2020 (MO20) (Figure 1A and Table S1). To compare heterologous adaptive immune responses, a second cohort of pigs was infected with H1N1 A/Missouri/CS20N08/2020 (MO20) virus two weeks after an initial infection with H1N1 A/California/04/2009 (pdmH1N1) and necropsied 5 days later (2X). While 1X pigs presented an increase in body temperature and shed more virus compared to 2X pigs, there was no difference in how the different genotypes responded to infection (Figure S1). Single-cell sequencing was performed on 12 pigs (3 per group), totaling 45,850 cells. A dimensionality reduction analysis identified 28 clusters by Uniform Manifold Approximation and Projection (UMAP) that were annotated according to established lineage markers (Figure 1B and Figure S2). The cell types most impacted by the number of times pigs were infected were CD8^+^ tissue resident memory T cells (TRM – cluster 4), cytotoxic CD8^+^ T cells (cluster 5), and cycling T cells (cluster 16) that were higher in 2X than 1X pigs, and CD2^-^ γδ T cells (cluster 9), and a subset of natural killer cells (NK2) cells (cluster 11) that were higher in 1X than 2X pigs (Figure 1C). The proportions of most cell types were comparable between genotypes with the exception that 2X *CD1D−/−* pigs had more CD4^+^ TRM cells (cluster 3) and fewer cytotoxic CD8^+^ T cells (cluster 5) after two infections compared to 2X *CD1D−/+* pigs. Next, we compared differentially expressed genes within individual cell types between 1X and 2X pigs by genotype (Figure 1D) or between *CD1D−/+* and *CD1D−/−* pigs by number of infections (Figure 1E). We found similar numbers of differentially expressed genes between *CD1D−/−* and *CD1D−/+* pigs and between 1X and 2X pigs. However, a cluster of monocytes and macrophages (cluster 17) from 2X *CD1D−/+* pigs had more upregulated genes than 2X *CD1D−/−* pigs. An ingenuity pathway analysis of canonical cellular immune response networks identified that several of these upregulated genes fell within “Interferon Signaling” (*IFI35, IFI6, IFIT1, IFNAR2, ISG15, MX1, PSMB8, STAT1, TAP1*) and “Antigen Presentation Pathway” (*CD74, HLA-DRA, PDIA3, PSMB8, PSMB9, TAP1*) pathways, suggesting that stimuli originating from NKT cells may have altered the maturation of these antigen presenting cells.

**Figure 1.**
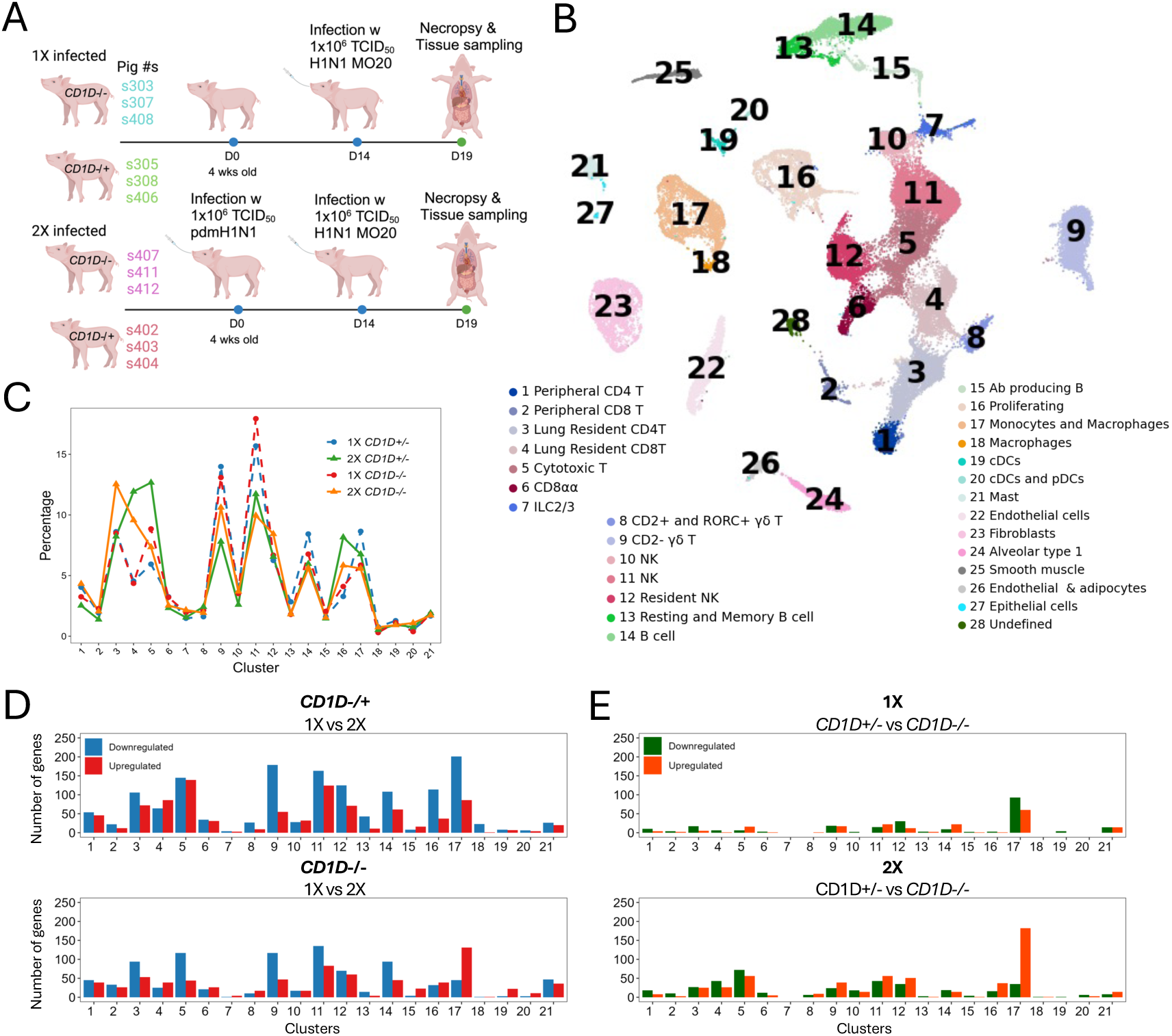
Single-cell transcriptomic analysis of IAV-infected *CD1D−/−* and *CD1D−/+* pig lungs. (A) Overview of experiment setup. 3 *CD1D−/−* and 3 *CD1D−/+* pigs were infected with pdmH1N1 IAV. Fourteen days later, the same 6 pigs were infected with H1N1 A/Missouri/CS20N08/2020 (MO20) IAV (designated 2X pigs) along with an additional 3 *CD1D−/−* and 3 *CD1D−/+* pigs (designated 1X pigs). Necropsies were performed 5 days after the MO20 infection to collect lung tissue for single-cell immune profiling. Created with BioRender. (B) Uniform manifold approximation and projection (UMAP) visualization of lung leukocyte populations colored by cell clusters. Clusters were identified using the graph-based Louvain algorithm at a resolution of 0.5. (C) The frequency of each cell type is presented for each treatment. (D) Bar graphs displaying the number of upregulated and downregulated differentially expressed genes (DEGs) in 1X compared to 2X *CD1D−/+* and *CD1D−/−* pigs. (E) Bar graphs displaying the number of DEGs in *CD1D−/+* compared to *CD1D−/−* pigs after 1X or 2X infections.

### TCR repertoire of lung T lymphocytes

We analyzed the pig lung tissue cells using our pig-specific V(D)J primers designed for compatibility with the 10x Genomics Chromium Next GEM Single-cell 5’ kit (Table S2). Over the 12 samples, we obtained 927,503,456 sequence reads an average of 77,291,955 reads per library, Data file 1). A de novo assembly of raw sequencing reads produced unannotated porcine V(D)J contigs which were identified using the sequencing primers and thereafter aligned to cells in the gene expression clusters in Figure 1. Across all samples, approximately 60% and 70% of identified TRA or TRB contigs were respectively aligned to single cells (Figure 2A, Data file 1). Assembled VDJ sequences were blasted against the international ImMunoGeneTics (IMGT) germline TRBV, TRBD, TRBJ, and TRAJ databases (13). Approximately 70% of TRB contigs in a pairing with TRA contigs mapped to an annotated TRVB or TRJB gene (Data file 1). Because Vα genes are not annotated in IMGT, Vα sequences were assigned according to pig TRAV sequences from our previous publication (14) (Figure S3, Data file 1). Only cells with IMGT-annotated TRB genes that were paired with productive TRA were used for further analysis (Figure 2B). These cells accounted for between 23% and 38% of cells within αβ T cell clusters (clusters 1-6) across samples.

**Figure 2.**
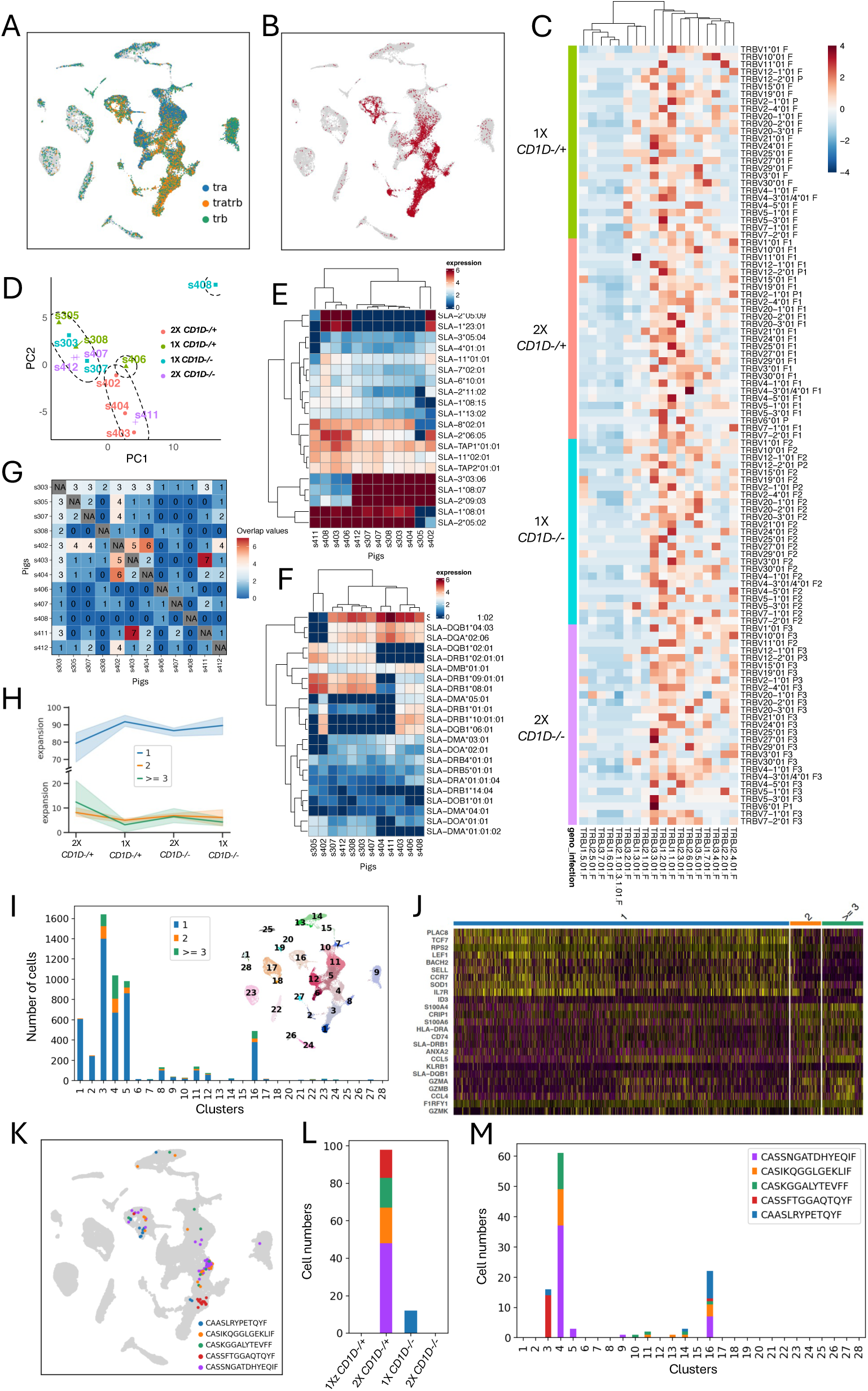
Characterization of T cell clonotypes from lung tissue. (A) UMAP plot of cells expressing one (tra, trb) or both (tratrb) TCR α and TCR β chains. (B) UMAP plot of cells with paired TCRα and TCRβ chains that were used for downstream analysis. (C) Relationship between TRBV and TRBJ usage in T cell receptor rearrangements by treatment. Cell barcode counts for TRBV and TRBJ gene segments were normalized by cell numbers across treatments and scaled by TRBV segments. (D) Principal component analysis of TRBV and TRBJ gene usages by individual pig. (E and F) Heatmaps showing row-scaled mean expression of the 10 highest differentially expressed SLA class I (E) and SLA class II (F) genes per pig. (G) Heatmap of overlapping clonotypes between pigs. (H) Proportion of clonotypes by treatment with 1, 2, ≥ 3 cells per clonotype. (I) Abundance of clonotypes by cluster with 1, 2, ≥ 3 cells per clonotype in the combined dataset. (J) Expression of naive and activation T cell markers in expanded and non-expanded clonotypes in the combined dataset. (K) UMAP plot displaying the five most expanded clonotypes defined by identical CDR3β region sequences in the combined dataset. (L and M) Number of cells in each of the five most expanded clonotypes by treatment (L) and cluster (M).

The expression of Vβ/Jβ and Vα/Jα combinations was analyzed to determine if favored rearrangements were correlated with *CD1D* genotype or the number of IAV infections (Figure 2C and Figure S3). Overall, *TRBJ3-3*01*, *TRBJ1-2*01*, and *TRBJ1-1*01* were used in many rearrangements, while pairings with *TRBJ1-5*01*, *TRBJ2-5*01*, *TRBJ3-7*01*, *TRBJ1-6*01*, and *TRBJ2-1*01* were relatively rare (Figure 2C). This is similar to our previous analysis of pig peripheral blood T cells that used a bulk RNA sequencing approach (14). We observed that 2X *CD1D−/+* pigs had Vβ/Jβ recombinations that were more similar to each other than to other pigs (Figure 2D). This was also the case for pigs s303, s305, s307, and s308, which were from the same litter. Since structurally rearranged TCRs are selected by major histocompatibility complex (MHC) molecules, we analyzed swine leukocyte antigen (SLA) class I and II expression in individual pigs by aligning scRNAseq transcripts across clusters (Figure S4) to the IPD-MHC database that provides a repository of MHC sequences for a number of species, including swine (13). To visualize SLA usage, a row-scaled mean expression of the 10 highest class I (Figure 2E) and class II (Figure 2F) genes in each pig was performed. Taking the expression of both SLA classes together, we could group pigs into 6 distinct MHC haplotype inheritance patterns (designated A-F) as follows: A – s403, s406, s408; B – s303, s307, s308, s407, s412; C – s305; D – s402; E – s411; F – s404. Vβ/Jβ pairings in the five B inheritance pattern pigs clustered together (Figure 2D), suggesting a link between MHC inheritance and VDJ selection.

Next, since TRB gene segments are fully annotated in IMGT whereas TRAV segments are not, we used identical CDR3β region sequences to analyze our samples for expanded clonotypes (Data file 1). Some clones were found in more than one pig, with the highest numbers of shared clones among the 2X infected pigs (Figure 2G). Additionally, 2X pigs, especially the *CD1D−/+* group, had more expanded clones than the 1X groups (Figure 2H). The highest concentrations of clonally expanded T cells were among CD4^+^ and CD8^+^ TRMs, cytotoxic CD8^+^ T cells, and cycling T cells while comparatively few expanded clones were present among peripheral-derived T cells in clusters 1 and 2 (Figure 2I). Expanded clones were enriched for immune activation/effector related genes compared to unexpanded clones (Figure 2J). 2X *CD1D−/+* pigs contributed four of the five most expanded clonotypes, all of which included more than twelve cells (Figure 2K and 2L). One of these clonotypes originated from CD4^+^ TRMs while the remaining four were from CD8^+^ TRMs (Figure 2M).

### BCR repertoire of lung B lymphocytes

Using a similar approach to our TCR profiling, we analyzed the pig lung tissue cells using our pig-specific primers for the IGK, IGL, IGHM, IGHA, and IGHG genes (Table S2), which respectively encode the immunoglobulin κ and λ light chains and IgM, IgA, and IgG heavy chains. Over the 12 samples, we obtained 1,395,728,419 sequence reads an average of 11,631,0701 reads per library (Figure 3A, Data file 1). Assembled V(D)J sequences were blasted against the IMGT germline IGL, IGK, and IGH databases (13). Only cells with IMGT-annotated IGH genes that were paired with productive IGL or IGK chains were used for further analysis. The percentage of B cells expressing different light and heavy chain contigs are as follows, 21% IGK, 40% IGL, 52% IGHM, 5% IGHG, and 3% IGHA. These cells accounted for between 53% and 70% of B cells among the 12 samples.

**Figure 3.**
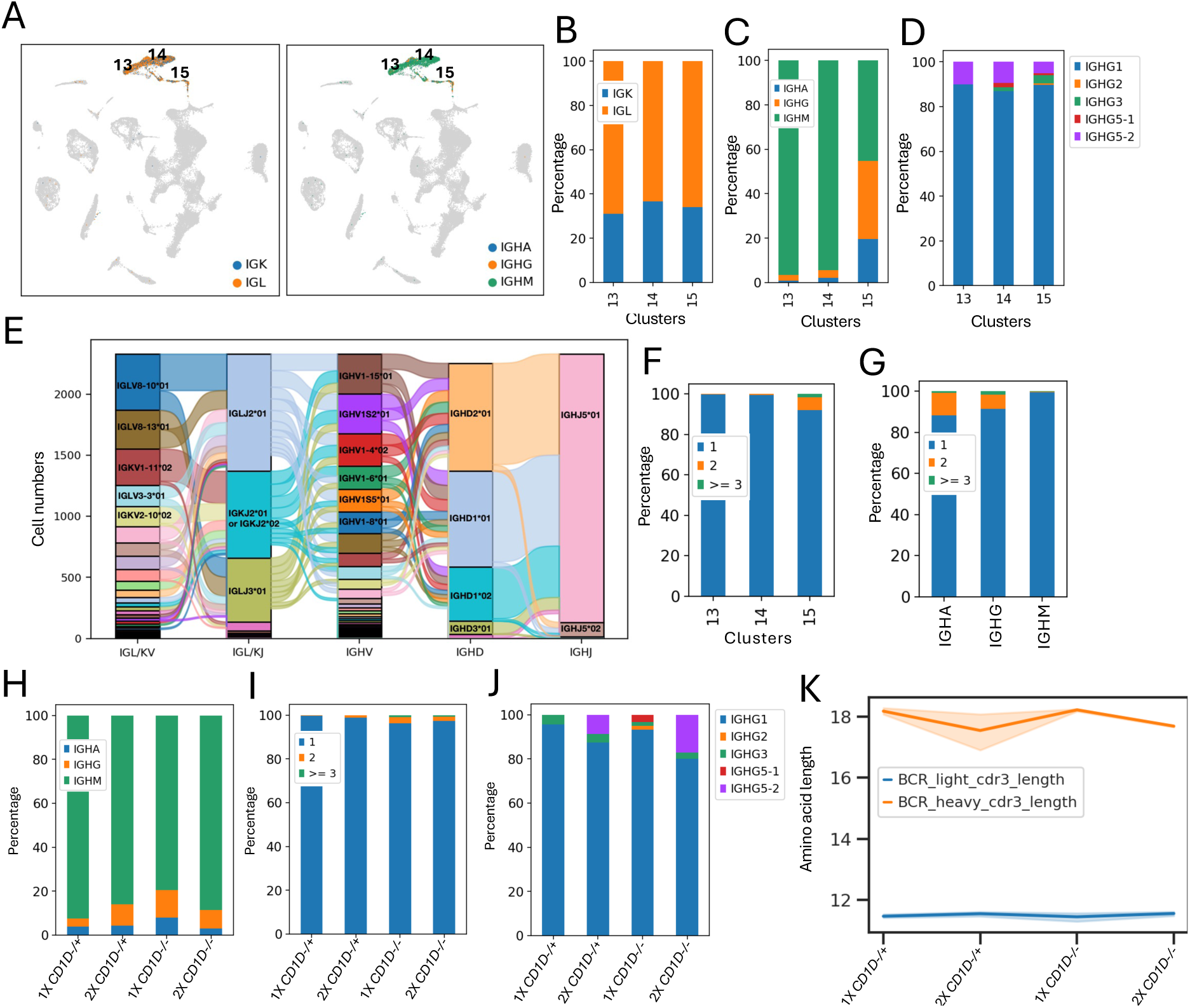
B cell receptor repertoire profiling of lung tissue B cells. (A) UMAP plots of cells expressing IGK and IGL chains and IGHM, IGHG, and IGHA chains. (B) Proportion of cells expressing IGK and IGL in each B cell cluster. (C) Percentage of cells expressing IGHM, IGHG, and IGHA in each B cell cluster. (D) Percentage of IGHG+ cells expressing different porcine IGHG subclasses in each B cell cluster. (E) Number of cells expressing light and heavy chain V(D)J gene segments. (F) Percentage of clonotypes by cluster with 1, 2, ≥ 3 cells per clonotype. (G) Percentage of clonotypes by IGH chain with 1, 2, ≥ 3 cells per clonotype. (H) Proportion of cells expressing IGHM, IGHG, and IGHA by treatment. (I) Percentage of clonotypes by treatment with 1, 2, ≥ 3 cells per clonotype. (J) Percentage of IGHG+ cells expressing different porcine IGHG subclasses by treatment. (K) Length of light and heavy chain CDR3 sequences by treatment.

The overall IGL to IGK ratio was 1.8 (Figure 3B). Prior studies have reported that IGL:IGK usage in circulating immature pig B cells is ∼1:1(15–17). Plasma B cells (cluster 15) had the highest proportions of IGHG+ and IGHA+ cells consistent with B cells that have undergone diversification for mucosal antibody secretion (Figure 3C). Since our IGHG primers targeted a conserved constant region of the eight genomic IgG constant region gene sequences cataloged in IMGT (*IGHG1, IGHG2, IGHG3, IGHG4, IGHG5-1, IGHG5-2, IGHG6-1, IGHG6-2*) (18–20), we mapped IGHG sequences to the IMGT database. *IGHG1* dominated all three clusters (Figure 3D). *IGHG5-2* was also detected in every B cell subset. However, only antigen-experienced memory B cells and plasma cells in clusters 14 and 15 exhibited *IGHG2*, *IGHG3*, and *IGHG5-1* subclasses. *IGHG4* and *IGHG6* could not be distinguished from the other subclasses.

Next, we analyzed V(D)J gene segment usage (Figure 3E, Figure S5 and Figure S6). Unlike humans or laboratory rodents, pigs use a limited number of IGHV, IGHD, and IGHJ genes to form most of their BCR repertoire, with >90% of the VDJ repertoire made up of seven IGHV genes, two IGHD segments, and a single IGHJ segment (21–23). Consistent with these studies, we found six IGHV genes (*IGHV1-15, IGHV1S2, IGHV1-4, IGHV1-8, IGHV1S5, IGHV1-6*) accounted for ∼70% of IGHV usage (Figure 3E and Figure S6) and that IGHD and IGHJ usage was largely restricted to *IGHD1* or *IGHD2* and *IGHJ5*.

We analyzed our samples for expanded clonotypes defined by the recombination of identical IG light and heavy chain CDR3 sequences (Data file 1). This revealed far fewer expanded B cells compared to the T cell compartment, with no single clonotype containing more than four cells. Almost all expanded clonotypes were among the small number of IgG or IgA secreting plasma cells (cluster 15) (Figure 3F and 3G). No striking differences in IGH usage (Figure 3H) and clonotype expansion (Figure 3I) were observed among the four treatment groups. However, the ratio of *IGHG5-2*/*IGHG1* was greater in 2X than 1X pigs (Figure 3J) while heavy chain CDR3 length tended to be greater in 1X than 2X pigs (Figure 3K).

### Longitudinal assessment of T cells from lung lavage fluid

Clonotype tracking is a powerful approach to monitor changes in the frequency of clonotypes of interest in cancer and vaccine immunology within a single individual (24–28). Using our swine TCR α and β chain primers, we analyzed T cells for V(D)J clonotypes and TCR α and β chain gene usage from the lungs of three specific pathogen free raised piglets (Figure 4A). Two pigs (FLU1 and FLU2) were vaccinated with a combination of inactivated pdmH1N virus and adjuvant at 28 days of age and boosted 15 days later. Both pigs were infected with live pdmH1N virus 2 weeks after the booster. Lung fluid was collected 3 days before infection (T1) and 7 days after infection (T2). The third pig (NAIVE) was lavaged at 28 (T1) and 33 (T2) days of age. Between 1,041 and 6,789 (average = 3,472) CD3^+^ cells sorted from each lung fluid sample (Figure 4B) were subjected to paired single-cell RNA and TCR sequencing. The combined dataset separated into 15 clusters (Figure 4C and 4D). Clusters 10 and 11 were γδ T cells that are relatively abundant in pigs. Clusters 1, 2, 3, 4, 5, 6, 7, 11, 12, 13, 14 presented tissue residency markers, while clusters 8, 9, and 10 expressed circulating T cell markers (Figure 4E). Tissue resident memory T cells were composed of both CD4^+^ (cluster 1) and CD8^+^ cells (clusters 2, 4, 5, 6, 12) as well as proliferating cells that contained a mixture of CD4^+^ and CD8^+^ TRMs (cluster 13, 14). In response to IAV infection, both FLU1 and FLU2 pigs presented an increase in the frequency of proliferating CD4^+^ and CD8^+^ TRMs, CD2^-^ γδ T cells, CD4^+^ TRMs, peripheral CD4^+^ T cells, naïve CD8αα T cells, and CD8^+^ tissue effector memory (TEM) clusters, and a decrease in the frequency of CD8^+^ TRMs (Figure 4F). We identified 10 modules of co-regulated genes and regulatory networks across cell types (Figures 4G and 4H, Data file 1) among which Module 5 in clusters 7, 8, 9, 10 harbored naïve/circulating T cell genes, Module 6 in clusters 2, 3, 4, 5, 6, 11 harbored tissue residency and cytotoxic genes, and Module 10 in clusters 1, 2, 11 harbored interferon stimulated genes (ISG).

**Figure 4.**
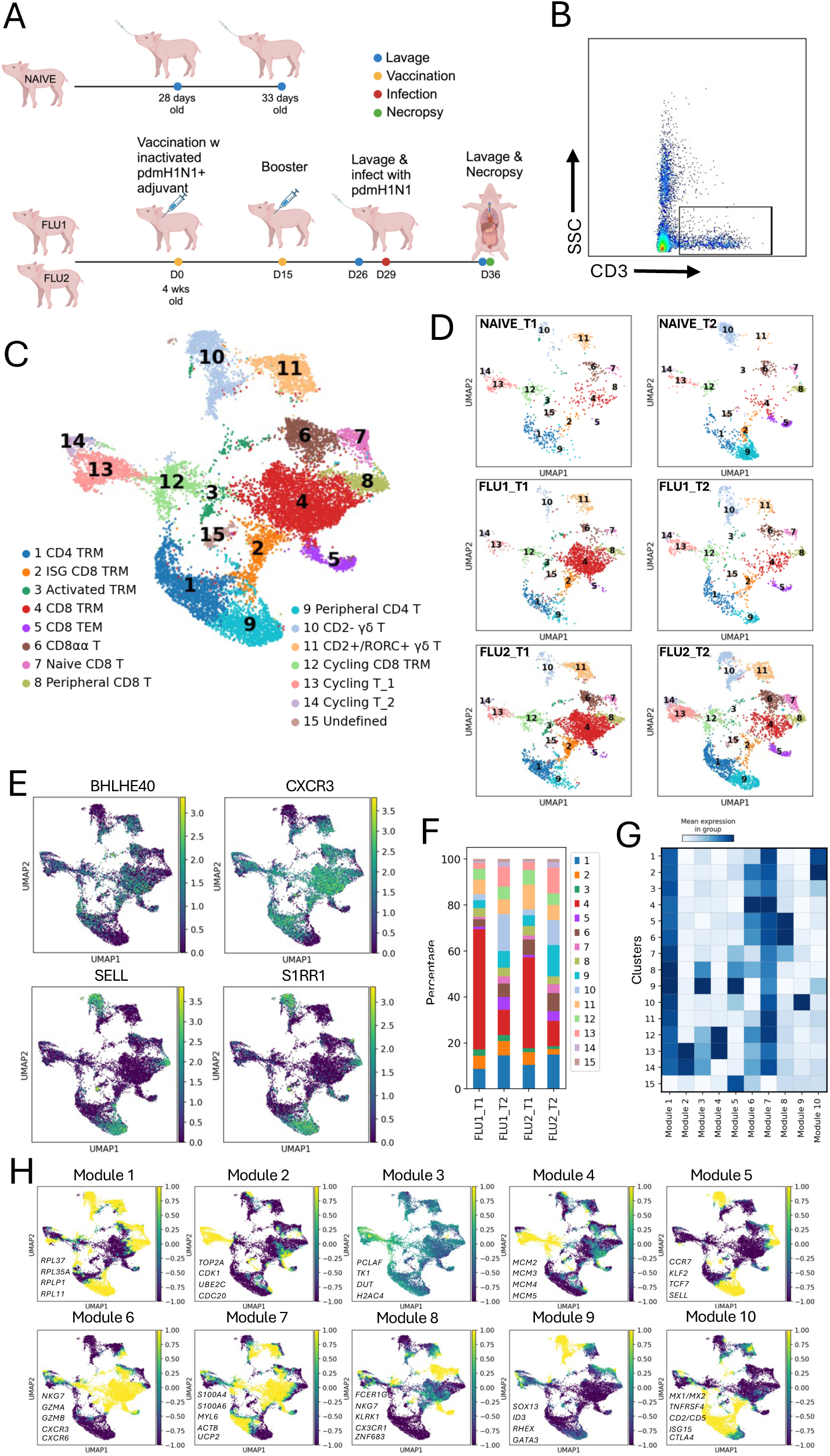
Single-cell transcriptomic analysis of T cells isolated from lung lavage fluid. (A) Overview of experiment setup. T cells were FACS-sorted from the lung lavage fluid of 3 infant pigs: two pdmH1N1-vaccinated and -infected pigs (FLU1 and FLU2) and one naïve pig (NAÏVE). FLU pigs were sampled 3 days before IAV infection (T1) and again 7 days after infection (T2). The NAÏVE pig was sampled at 28 (T1) and 33 (T2) days of age. Created with BioRender.. (B) FACS plot showing acquisition of lung lavage T cells. (C) UMAP plot of the combined T cell datasets. (D) UMAP plots displaying individual samples. (E) Examples of genes used to identify resident (*BHLHE40, CXCR3*) and circulating (*SELL, S1PR1*) T cells. (F) Proportions of T cell subsets in FLU1 and FLU2 pigs at T1 and T2 timepoints. (G) Heatmap of 10 gene modules whose genes had a similar expression pattern across cell clusters. (H) UMAPs showing select genes from modules 1-10.

Approximately 60% of αβ T cells recovered had paired TCRα and TCRβ chains (Figure 5A). Sequence identity can be used to map the phylogenetic relatedness of TCRα and TCRβ chains in individual samples (Figure 5B). Many clones, identified by CDR3β sequences, were present at both T1 and T2 in the same animal, especially the FLU2 pig (Figure 5C). A high proportion of expanded clones were CD8^+^ TRMs (Figure 5D). The two vaccinated and infected pigs presented the five most expanded clones, especially the FLU2 pig (Figure 5E and 5F). The low number of expanded clonotypes present in the NAÏVE pig samples is due partly to the lower number of cells we were able to collect from this pig. Next, we examined a larger collection of expanded clonotypes in FLU1 and FLU2 pigs, before and after infection, to identify potential influenza-reactive T cells (Figure 5G). While the frequency of several clones decreased from T1 to T2, some increased, including CSAGERSNYEQIF, CASSVRSYPLNDLHF, CASSFGGVHTGQLYF, CAWSTTGTVTGQLYF, and CSAGEGGFGDTCFF. We also analyzed specificity groups within the CDR3β repertoire using immunarch (29, 30), a program that enables clustering of TCRs with an increased probability of sharing antigen specificity due to conserved motifs of CDR3 sequences. Two CDR3β 4-mer motif patterns (ASSL and SSLV) were found to be enriched in FLU1 and FLU2 pig samples (Figure 5H). Interestingly, a few of the CDR3β sequences in expanded clones were identical or differed by a single amino acid from curated human TCR CDR3β sequences which recognize IAV epitopes in M1, NP, PB1, and PB2 (31). For example, the seventh most expanded CDR3β sequences in all pigs, ASSPGQGYEQ, matches a human CDR3β sequence that recognizes the immunodominant IAV Matrix protein 1 epitope GILGFVFTL when presented by human HLA-A*0201 (Figure 5I) (32, 33). This may arise because peptide binding motifs of some common swine SLA molecules partly overlap with the binding motifs of human HLA molecules, including a number of HLA-A*0201-restricted IAV peptides (34).

**Figure 5.**
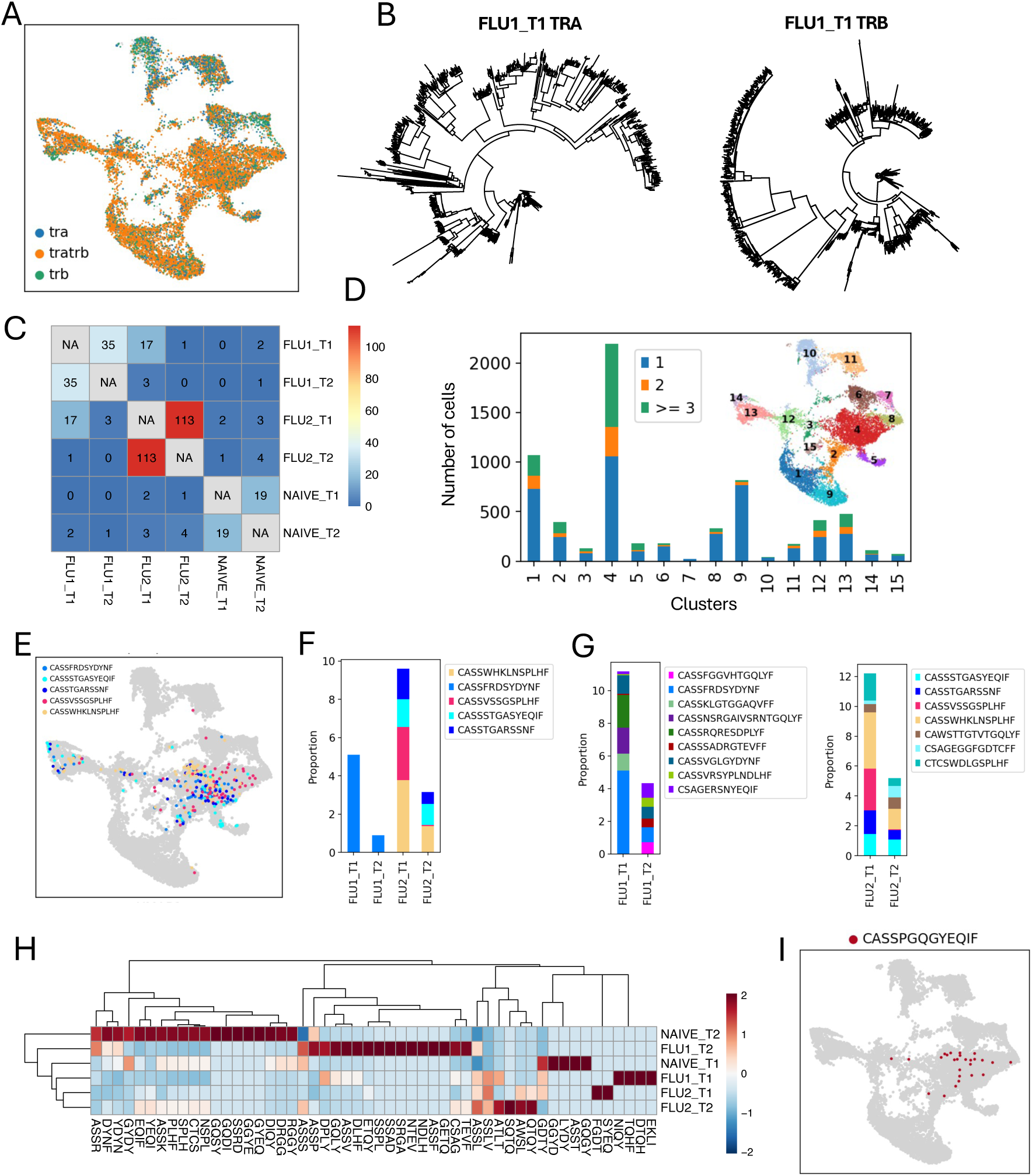
T cell clonotype tracking using VDJ recombination at the TRB locus. (A) UMAP plot of cells expressing one (tra, trb) or both (tratrb) TCR α and TCR β chains in the combined dataset. (B) Phylogenetic trees of TCRα and TCRβ sequences in the FLU1_T1 sample. (C) Heatmap of overlapping clonotypes between samples. (D) Abundance of clonotypes by cluster with 1, 2, ≥ 3 cells per clonotype in the combined dataset. (E) UMAP plot displaying the five most expanded clonotypes in the combined dataset. (F) Number of cells in each of the five most expanded clonotypes by sample. (G) Proportion of the most abundant clonotypes in FLU1 and FLU2 pigs by sample time. (H) Heatmap displaying the most abundant CDR3β 4-mer motifs in each pig normalized by cell numbers in each sample. (I) T cell clones expressing the CDR3β sequence ASSPGQGYEQ that matches a human CDR3β sequence recognizing the human HLA-A*0201 M1 epitope GILGFVFTL.

## Discussion

The current study describes assays for single-cell TCR/BCR sequencing in pigs which presents a useful tool for enhancing effective vaccine and therapeutic design to protect swine health and to increase the potential of pigs as biomedical models for studying human immune-related physiological processes and diseases. We examined the utility of the assay in the setting of the pulmonary immune response against IAV infection since influenza is a respiratory pathogen of major importance for both humans and swine (35). One set of samples were from cryopreserved lung sections of *CD1D*-expressing and -deficient pigs after one IAV infection or two infections with heterologous IAVs. This is of interest since NKT cell effector responses are important for anti-IAV immunity in mice, including that NKT cell-deficient mouse strains are significantly more susceptible to IAV infections than standard mice (36–39). Moreover, NKT cells are capable of a wide array of CD4 T helper cell functions, which have the capacity to elicit wide-ranging cellular and humoral responses that can substantially boost the quality and durability of immune responses, including against heterologous and heterosubtypic IAV infections (40). Among the 12 samples in this dataset, we detected almost all VDJ gene segments annotated for TCR β chains, including some that were annotated as pseudogenes. In addition, most of the Vα genes detected overlapped with TCR α chains that we identified in a previous analysis of TCR chain usage using bulk RNAseq (14).

While we found that cells which expressed both α and β TCR chains were mostly within αβ T cell clusters (clusters 1, 2, 3, 4, 5), we also detected cells expressing unpaired α and β chains in some non-αβ T cell clusters, such as CD2^-^ γδ T cells that expressed TCR β chains and NK cells and B cells that expressed TCR α chains. This is consistent with prior reports that a significant proportion of peripheral γδ T cells express TCR β (41) chains and that NK cells express germline TCR transcripts (42). The NK cell observation is supported by the fact that virtually all NK cell TRA transcripts matched to TRAJ segments but very few to TRAV segments. Examination of Vβ/Jβ combinations confirmed previous studies showing that certain rearrangements are favored in pigs (14, 43). In several pigs, VDJ rearrangements clustered by MHC inheritance pattern. We also found that VDJ rearrangements preferred by 2X *CD1D−/+* pig clonotypes clustered together which could mean that IAV infection influenced Vβ/Jβ selection in NKT cell-intact pigs.

The most expanded clones were among CD4^+^ and CD8^+^ TRMs, cytotoxic CD8^+^ T cells, and proliferating T cells, consistent with reports that these populations harbor antigen experienced T cells that are poised for rapid responses during an IAV infection (44). As expected, 2X pigs had more expanded clones than 1X pigs. This was less apparent in *CD1D−/−* compared to *CD1D−/+* pigs, which might suggest that induction of IAV-specific lung T cells is reduced in the absence of NKT cell helper functions.

Lung tissue B cells consisted of naïve B cells, memory B cells, and a small population of plasma cells. Plasma cells displayed the highest diversity in immunoglobulin heavy chain usage and clonally expanded B cells. Consistent with prior reports (15–18, 20, 21, 23, 45, 46), we found (i) a mixture of IGL+ and IGK+ B cells, (ii) IgG heavy chain usage was dominated by the IGHG1 subclass, and (iii) a limited number of IGHV, IGHD, and IGHJ genes formed most of the BCR repertoire. Pigs are interesting in so far as their pathway of antibody repertoire development has evolved somewhat differently from mice and humans, including that they possess a highly streamlined IGH gene complex, which contains IGHV genes that all belong to a single ancestral IGHV3 family, and that only one IGHJ segment is functional (18, 20, 21, 23, 45). Because of the small number of IGHV, IGHD, and IGHJ segments used, the combinatorial diversity in pigs is comprised of a mere ∼14 possibilities compared to ∼9,000 in humans (23).

TCR transcript capture was more efficient in longitudinally collected lavage fluid T cell samples than in the *CD1D* lung tissue samples, probably owing to the former consisting entirely of FACS-purified T cells. As in lung tissue cells, most αβ T cell clusters exhibited paired α and β TCR chains whereas a significant fraction of CD2^-^ γδ T cells expressed TCR β chains and CD2^+^ γδ T cells and CD8αα T cells were enriched for TCR α chains. Clusters with the highest numbers of expanded clones were again tissue resident memory T cell populations. Our ability to detect a substantial number of the same T cell clones at two different timepoints in the same individual shows the utility of TCR profiling for monitoring changes in clonotypes of interest in pigs. The higher number of expanded clonotypes within FLU1 and FLU2 pig samples, some of which overlapped, compared to the NAÏVE pig, may be the result of IAV exposure by vaccination and infection leading to an increase in antigen experienced T cells in lung tissue and/or because fewer cells were collected in the NAÏVE pig. While there is a lack of reagents to identify influenza-specific T cell clones in commercial pig breeds, this information can to some extent be inferred by studying clonotypes in vaccinated pigs that increase after virus exposure. Using this approach, we identified CDR3 sequences that are potentially reactive to IAV antigens, including a clone with a CDR3β sequence identical to a human motif that recognizes an immunodominant epitope from Matrix protein 1 (34).

In summary, the assays presented in this study can easily be applied to 5’ 10x Genomics protocols for use in swine. Our protocols can be employed to profile TCRs and BCRs in the same sample which enhances the utility of the method as most adaptive immune responses involve both cellular and humoral responses. Accordingly, the combined protocol could shed light on acquired immunity that develops in response to vaccination and infection in production pigs, as well as in the growing number of immune-related pig models being developed for biomedical use.

## Methods

### Pigs

The National Swine Resource and Research Center (NSRRC) at the University of Missouri bred a boar homozygous for a *CD1D* gene deletion with two sows heterozygous for the *CD1D* deletion that were full sisters to produce piglets that were homozygous a the *CD1D* deletion (*CD1D−/−*) as well as heterozygous segregants (*CD1D−/+*). Our previously described *CD1D* breeding herd (47) is on a commercial Large White crossbred background and maintained under specific pathogen free conditions. The *CD1D* genotypes of pigs were determined by PCR and flow cytometry as previously described (47, 48). Piglets used for longitudinal assessment of T cells from lung lavage fluid were commercial Large White crossbred background pigs provided by the NSRRC.

### Virus infection and sample collection

Eight *CD1D−/−* and eight *CD1D−/+* pigs were transferred to biocontainment rooms at 4 weeks of age after being confirmed seronegative for IAV nucleoprotein antibodies by ELISA developed by at the Veterinary Diagnostic Laboratory at the Iowa State University. At day 0, 4 *CD1D−/−* and 3 *CD1D−/+* pigs were intratracheally infected with 1×10^6^ tissue culture infectious dose (TCID_50_) of H1N1 A/California/04/2009 (pdmH1N1) in 2mL of DMEM (Gibco, Brooklyn, NY) after sedation with a combination of midazolam, butorphanol, and xylazine. Fourteen days later, all 16 pigs were sedated and infected with 1×10^6^ TCID_50_ of H1N1 A/Missouri/CS20N08/2020 (MO20) IAV. Pigs were measured for clinical disease signs and daily nasal swab virus titers as previously described (49). Five days later, all pigs were sedated and euthanized by pentobarbital sodium intracardiac injections (70 mg/kg of body weight). At necropsy, the lungs were removed from the thoracic cavity for tissue collection. and euthanized. Cells were isolated from 3 animals of each genotype from single (1X) and twice (2X) infected pigs for scRNAseq and receptor profiling. Litter, sex, *CD1D* genotype, and MHC inheritance pattern of the piglets used for sequencing are described in (Table S1).

For the study to collect T cells through successive lung lavages, two pigs (FLU1 and FLU2) were intramuscularly vaccinated with a combination of 1×10^6^ TCID_50_ of ultraviolet-inactivated pdmH1N1 virus and an oil-in water adjuvant (Emulsigen, 1:5 vaccine volume) at 28 days of age and boosted 15 days later. Both pigs were intratracheally infected with 1×10^6^ TCID_50_ live pdmH1N1 virus 2 weeks after the booster. Lung fluid was collected 3 days before and 7 days after infection. A third unvaccinated, uninfected pig (NAÏVE) was lavaged at 28 and 33 days of age. Pigs were intratracheally sedated with midazolam, butorphanol, and xylazine to perform lung lavages and infections. Lavages involved inserting a size 10 French catheter attached to a syringe into the lung after which the lung was flushed twice with 5 ml of sterile saline solution. Recovered lung fluid was ejected into 10 ml phosphate buffered solution containing 10% fetal bovine serum (FBS).

The studies were in accordance with the University of Missouri’s Institutional Animal Care and Use Committee (protocol number 34343) and Institutional Biosafety Committee (protocol number 17320).

### Tissue sampling and cell isolation

Approximately 1 g of tissue collected from the left cranial, middle, and caudal lung lobes, were combined, and then digested with 2.5 mg/mL of Liberase TL (Roche, Indianapolis, IN) in Dulbecco’s Modified Eagle Medium (Thermo Fisher, Waltham, MA) at 37°C for 45 minutes. The digested tissue was dispersed into single cells as previously described (50), and then passed through a 70 μm cell strainer (Thermo Fisher, Waltham, MA). Cells were immediately cryopreserved in freezing media [90% FBS, 10% dimethylsulfoxide (DMSO)] in temperature-controlled freezing containers at 3 × 10^7^ cells per/mL and stored at −80°C until use. Samples were thawed in thawing media (RPMI –HyClone, Logan, UT, - 20% FBS), resuspended in RPMI with 10% FBS, filtered through a 40μm filter (Bel-Art SP Scienceware, Wayne, NJ), and counted for viable cells using a Countess 3 automated cell counter (Thermo Fisher, Waltham, MA). To obtain T cells from lung washes, lavage fluid was filtered first through a 70 μm cell strainer (Thermo Fisher, Waltham, MA) and then a 40 μm pipette tip Flowmi® cell strainer (SP Scienceware, Warminster, PA), washed in RPMI with 10 % FBS, counted, stained with PE-Cy7-conjugated anti-porcine CD3 antibody (clone BB23-8E6-8C8; BD Biosciences, San Jose, CA) and propidium iodide viability dye, and sorted for live CD3 positive cells using a BD FACSMelody Cell Sorter (BD Biosciences, San Jose, CA). Approximately 10,000 cells from cryopreserved lung tissue and between 1,041 and 6,789 T cells from lung lavage fluid were loaded onto the 10x Chromium controller (10x Genomics, Pleasanton, CA).

### Primer design

Pig-specific V(D)J primers were designed according to guidelines from the 10x Genomics Chromium Next GEM Single-cell 5’ V2 user guide, which is well described in a recent publication on scTCRseq in dogs(51). This protocol employs a nested PCR design which involves two rounds of V(D)J amplification using the same forward primer that primes off the 5’ Illumina adapter sequence which is annealed to the 10x barcoded cDNA during the conversion from mRNA. The first round of amplification uses the 5’ forward primer in combination with a 3’ outer reverse primer that matches the C region of the targeted chain. The second round uses the same forward primer in combination with a second reverse primer that primes the C region at an inner 5’ region from the outer reverse primer. To design the 3’ reverse primer sets, C-gene sequences were identified for TRA, TRB, IGH, and IGL and IGK transcripts. Inner and outer primers were respectively designed to target regions between 50-200 and 200-300 base pairs away from the 5’ region of the C region. Primer candidates were selected based on the percent of GC content, similar melting temperature range, and minimal predicted interaction with other regions in the pig genome. The final primer sequences as well as accession numbers for the DNA and rearranged mRNA transcripts used to identify the C regions are listed in (Table S2).

### Single cell processing

Libraries were constructed by following the manufacturer’s protocol with reagents supplied in the Chromium Next GEM Single Cell 5′ Kit v2 (10x Genomics). Briefly, cell suspension concentration and viability were measured with a Cellometer K2 (Nexcelom Biosciences, Lawrence, MA) stained with an acridine orange/propidium iodine dye mix (Invitrogen, Waltham, MA). Cell suspension combined with reverse transcription master mix were loaded on a Chromium Next GEM chip K along with gel beads and partitioning oil to generate gel emulsions (GEMs). GEMs were transferred to a PCR strip tube and reverse transcription performed on a Veriti thermal cycler (Applied Biosystems, Waltham, MA) at 53°C for 45 minutes. All samples underwent 11 cycles of cDNA amplification upon which cDNA concentration and quality were assessed using a Fragment Analyzer 5200 (Agilent, Santa Clara, CA). For the gene expression library, up to 50 ng of the cDNA was fragmentated, end-repaired, A-tail added, and ligation of sequencing adaptors was performed according to manufacturer specifications. V(D)J libraries were constructed using pig-specific primers for two successive enrichments of TCR and BCR transcripts. The TCR and BCR assay used primer pools containing 1.43 μM per gene specific primer and 1.43 μM of the 10x forward primer. Nested PCR amplification of the TCR and BCR sequences was performed using adapted mouse and human PCR protocols. This involved amplifying 2 μL of cDNA in 100 μL total reaction volume using 50 ul Amp Mix (10x Genomics). Steps involved in the first reaction were 98 °C for 45 s for initial denaturation followed by 9 cycles of 98 °C for 20 s, 65 °C for 30 s, and 72 °C for 60 s. PCR product was purified using AxyPrep MagPCR Clean-up beads (Axygen) and subjected to a second round of amplification using the same cycling conditions to the first except a total of 8 cycles was used. The amplicons were purified using AxyPrep MagPCR Clean-up beads (Axygen). Libraries were constructed from PCR amplicons according to manufacturer specifications. The concentrations for all libraries were measured with the Qubit HS DNA kit (Invitrogen) and fragment sizes were determined on a Fragment Analyzer 5200 (Agilent). Libraries were pooled and sequenced on a NovaSeq 6000 (Illumina, San Diego, CA) to obtain paired end reads. The minimum sequencing depths targeted were 40,000 reads per cell for 5’ GEX libraries and 10,000 reads per cell for V(D)J libraries.

### Single-cell RNA sequencing data analysis

The Sscrofa 11.1 genome assembly was used to align sequencing reads to generate gene matrix data by Cell Ranger (v8.0.0). Clustering analyses were performed using Seurat (v.4.4.0)(52). To filter out low-quality genes and cells, only genes expressed in more than 3 cells and cells with more than 200 genes and less than 10% mitochondrial reads were included in the analysis. Afterward, we followed a standard integration workflow to integrate samples. Briefly, transcript counts were log normalized, and the top 2,000 most variable genes in each dataset were identified using the *FindVariableFeatures* function. Then, the *SelectIntegrationFeatures* function was applied to genes that were consistently variable across datasets. Next, the *FindIntegrationAnchors* function identified a set of anchors between datasets using the top 30 dimensions from the canonical correlation analysis to specify the neighbor search space. Next, an integrated dataset was created by running the *IntegrateData* function. Then, clustering analysis workflow was performed using *RunPCA*, *FindNeighbours*, *FindClusters*, and *RunUMAP*. Cell types were assigned based on the expression of known cell type markers (Figure S2).

### Single-cell V(D)J data analysis

Single-cell TCR sequencing reads were assembled into contigs using cellranger vdj (10x Genomics) pipeline in denovo mode rather than reference-based mode due to the incomplete annotation of the pig germline Vα chain sequences. To identify the V(D)J chains, we searched assembled contigs against inner-enrichment primers using the usearch_global command. Primer matched TCR αβ and BCR IGH and IGL and IGK chains were selected and integrated with the above cellular gene expression profiles. The TCR β chains with matched TCR α chains were selected for downstream analysis. TRBV, TRBD, TRBJ, and CDR3 sequences were mapped to the pig TRB reference in IMGT using the IMGT/V-QUEST sequence alignment software. Cells with duplicated or multiple contigs were removed. Immunarch (v1.0.0) (29) was used to track clones across samples and identify k-mers from CDR3 sequencing. Scirpy (v.0.12.0) was used to analyze TCR β repertoires for clonal expansion (each unique CDR3) using the scirpy.pl.clonal_expansion command (53). In addition, CDR3 sequences were searched within the Immune Epitope Database (IEDB) (31) to find matches predicted to recognize epitope specificity using TCRMatch T cell epitope prediction tool. The unique contigs of TCR α and β chains in each cell were aligned to each other using CLUSTALW (https://www.genome.jp/tools-bin/clustalw), after which phylogenetic trees were generated to infer the similarities among contigs using the package ape (v5.7-1)(54). Using MMseqs2 (55), TRAJ sequences were mapped to the pig TRAJ reference in IMGT while TRAV segments were annotated according to TRAV sequences deposited in in GeneBank (55) that we previously named according to similar human TRAV genes (14). Similarly, BCR repertoires were annotated using the available pig IGH and IGK/IGL reference in IMGT using IMGT/V-QUEST(56). The reference mapped IGH chains and IGK/L chains were selected for downstream analysis. The IGH constant segments were mapped to IMGT/GENE-DB database. Scirpy was used to analyze clonal (the recombination of IGH and IGL CDR3s) expansion, gene usage, and CDR3 sequencing length analyses.

### SLA alleles annotation from scRNA-seq data

The Cellranger mkgtf was used to build the SLA alleles reference using the SLA FASTA files downloaded from the IPD-MHC database(13). Afterwards, SLA reads were quantified using Cellranger count command, SLA counts were log normalized, and then the average expression of each SLA allele in each sample was computed with the AverageExpression function in Seurat. The data were visualized in heatmap using ComplexHeatmap (v2.25.1) (57) and scaled for PCA analysis.

## Data availability

The sequencing data are available at Gene Expression Omnibus (accession GSE277475). Processed single-cell RNA sequencing objects are available for online visualization at https://singlecell.broadinstitute.org/single_cell/study/SCP2783 and https://singlecell.broadinstitute.org/single_cell/study/SCP2779. All relevant data are available from the authors.

