## Supplemental data for "Single-Cell Antigen Receptor Sequencing in Pigs with Influenza"

Table S1. Description of animals used in the study

| **Genotype** | Pig ID | # of infections | Gender | Litter | MHC Pattern |
|---|---|---|---|---|---|
| ***CD1D-/-*** | s303 | 1X | F | 1 | B |
| ***CD1D-/-*** | s307 | 1X | M | 1 | B |
| ***CD1D-/-*** | s408 | 1X | M | 2 | A |
| ***CD1D-/-*** | s407 | 2X | M | 2 | B |
| ***CD1D-/-*** | s411 | 2X | M | 2 | E |
| ***CD1D-/-*** | s412 | 2X | M | 2 | B |
| ***CD1D-/***+ | s305 | 1X | M | 1 | C |
| ***CD1D-/***+ | s308 | 1X | M | 1 | B |
| ***CD1D-/***+ | s406 | 1X | M | 2 | A |
| ***CD1D-/***+ | s402 | 2X | F | 2 | D |
| ***CD1D-/***+ | s403 | 2X | M | 2 | A |
| ***CD1D-/***+ | s404 | 2X | M | 2 | F |

Table S2. Porcine custom primer sets for scTCR/BCRseq

| **Primer name** | Target | 5'-3' Sequence | Accession | Ref. type |
|---|---|---|---|---|
| TCRa outer | TRAC | ATCGGTGCTTTTGCTCCAAG | MN086839.1 | mRNA |
| TCRa inner | TRAC | GTGGGCTCCGAGTCTTTTGT | MN086839.1 | mRNA |
| TCRb outer | TRBC | TCAGACAGTAGCTGGAGTCATTGAG | AB079894.1 | DNA |
| TCRb inner | TRBC | TCCGATGGTTCAAACACGGC | AB079894.1 | DNA |
| IgA outer | IGHAC | TGCACTTGGCACTTCAGGAT | AB699688.1 | DNA |
| IgA inner | IGHAC | CAATAACGCCCTCGCGACTA | AB699688.1 | DNA |
| IgG outer | IGHGC | CTGAGGGAGTAGAGCCCTGA | AB699686.1 | DNA |
| IgG inner | IGHGC | GCTCGGGGAAGTAGCTTGAG | AB699686.1 | DNA |
| IgM outer | IGHMC | AAGTACTTGCCGCCTCTCAG | AB699686.1 | DNA |
| IgM inner | IGHMC | GATGTTCTGGCTGCTGACCT | AB699686.1 | DNA |
| Kappa outer | IGKC | GAAGCTTTTGACCAGAGGGGA | KF561240.1 | mRNA |
| Kappa inner | IGKC | TCCAGGATGCCACTGCTTTG | KF561240.1 | mRNA |
| Lambda outer | IGLC | CGTCACTGTCTTCTCCACAATG | M59322.1 | mRNA |
| Lambda inner | IGLC | TCTGTTTCGAGGGCTTGGTG | M59322.1 | mRNA |

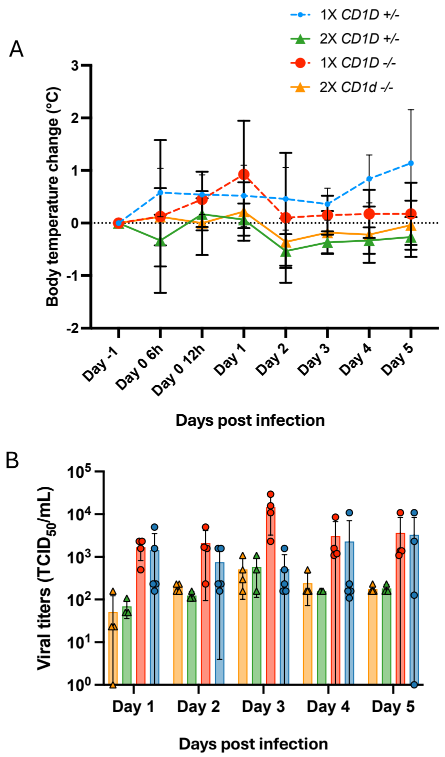


**Figure S1**. Results of experiment 1. (A) Change in body temperature during the challenge period based on body temperature at 0 d.p.i. (B) Viral titers in nasal swabs collected after challenge with H1N1 A/Missouri/CS20N08/2020.

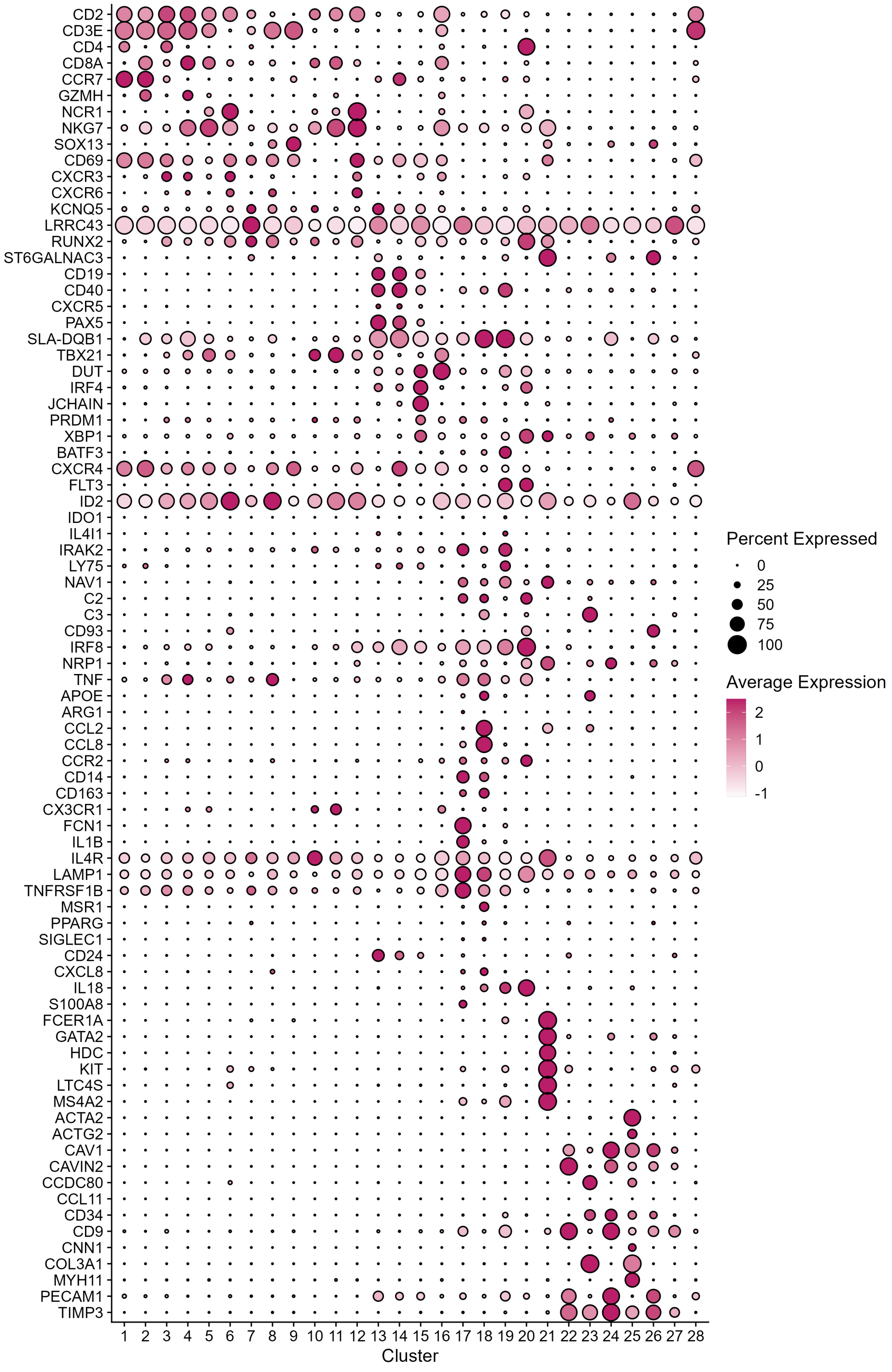


**Figure S2**. Dotplot showing row-scaled mean expression of some of the marker genes that were used to designate cell types to cell clusters in Figure 1B.

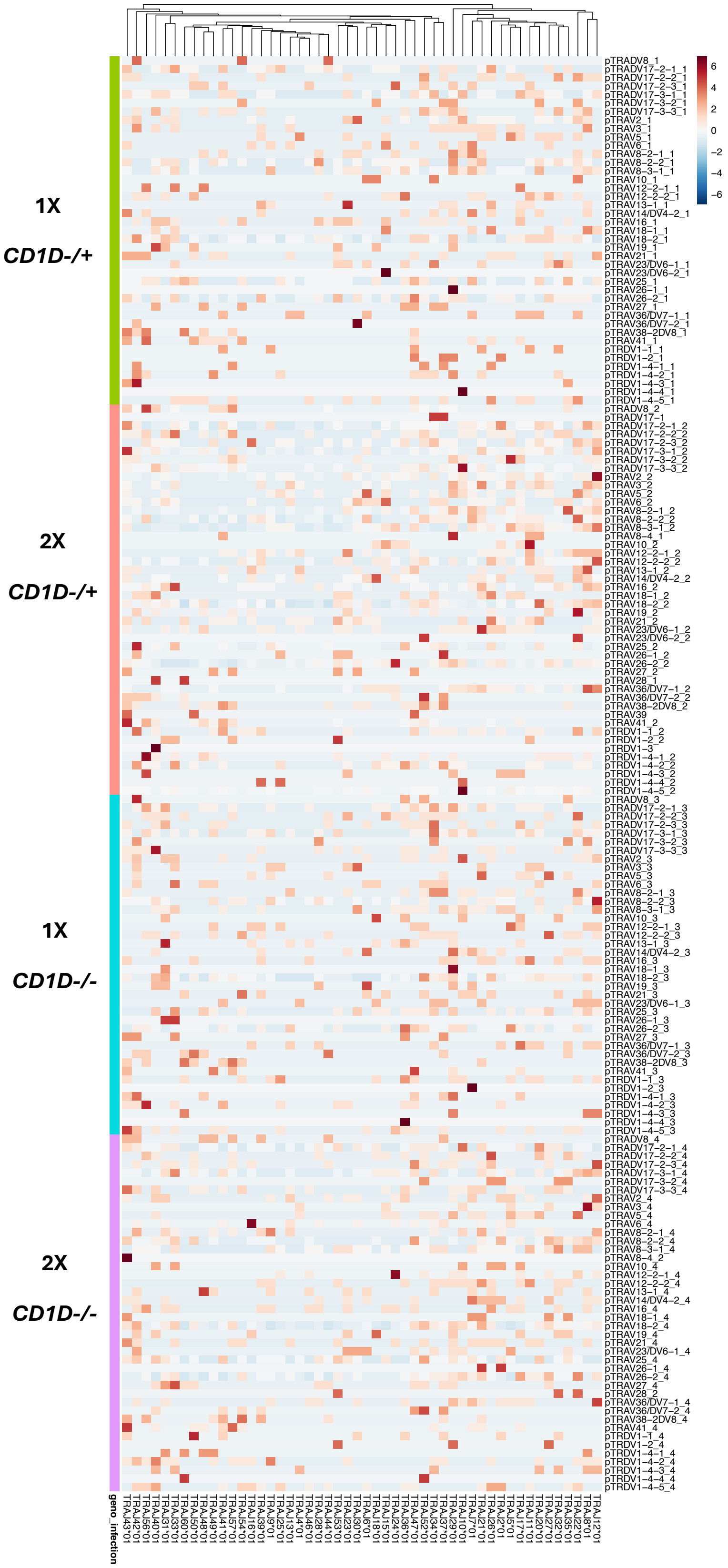


**Figure S3**. Relationship between TRAV and TRAJ usage in T cell receptor rearrangements by treatment. Cell barcode counts for TRAV and TRAJ gene segments were normalized by cell numbers across treatments and scaled by TRAV segments.

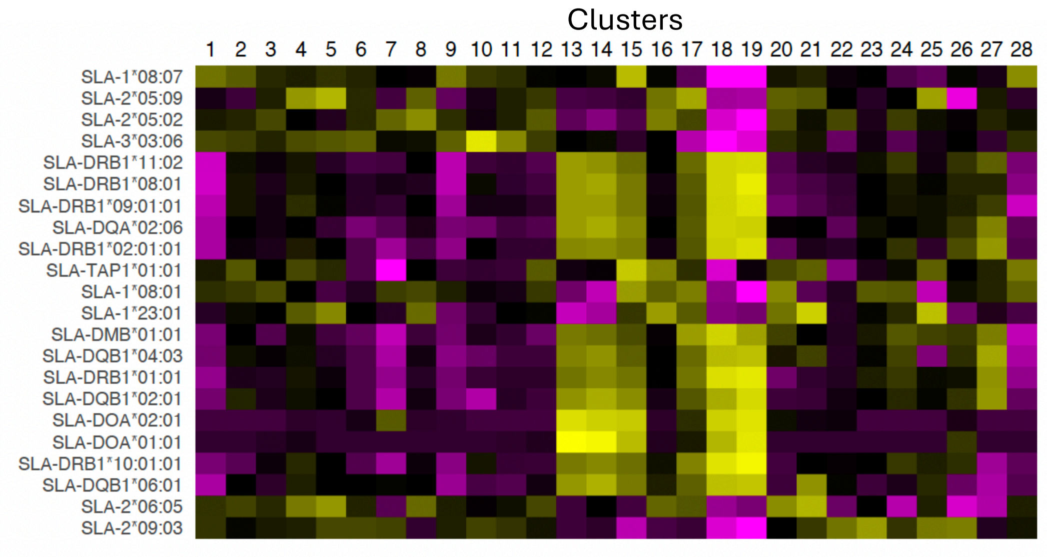


**Figure S4**. Heatmap showing row-scaled mean expression of SLA genes by cell type which was used to visualize SLA usage in each pig in Figures 2E and 2F.

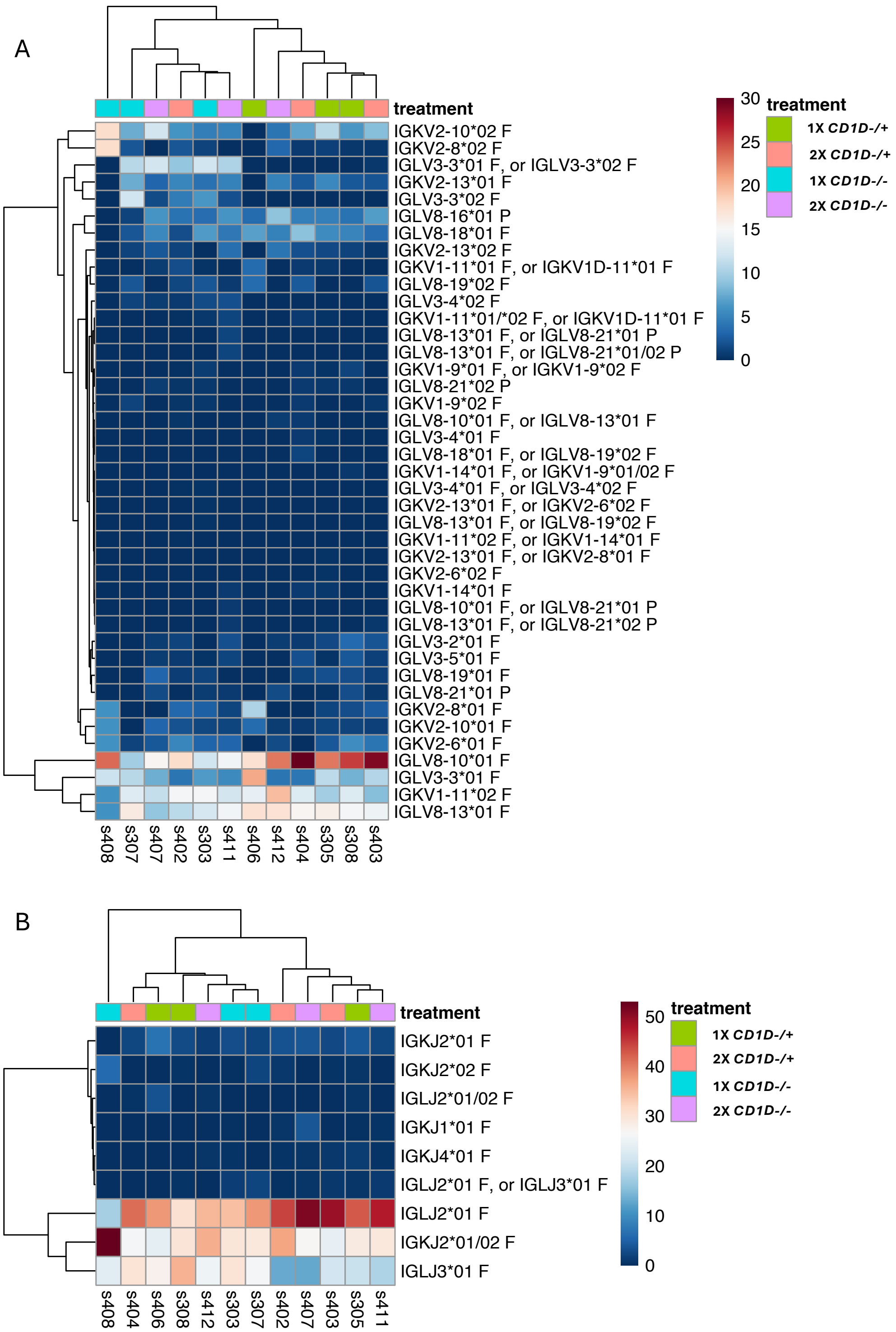


**Figure S5.** Heatmap showing row-scaled mean expression of V (A) and J (B) segment usage in LGL and LGK chains.

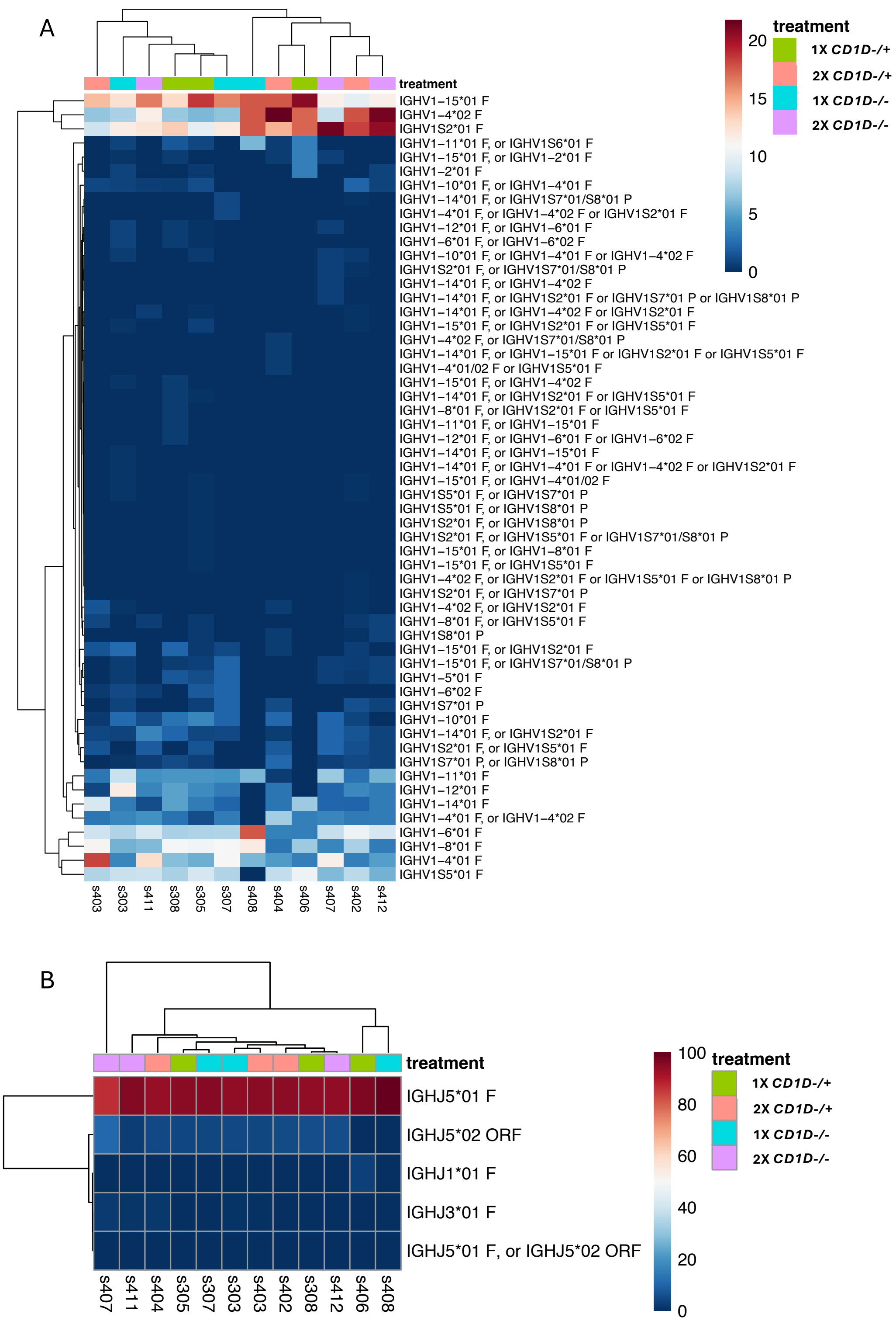


**Figure S6.** Heatmap showing row-scaled mean expression of V (A) and J (B) segment usage in IGHM, IGHA, and LGHG chains.
